# A phylogeny-guided framework for decoding mechanisms of human endogenous retrovirus regulation in health and disease

**DOI:** 10.64898/2026.05.19.726217

**Authors:** Andrew Patterson, Bryant Duong, Leena Yoon, Maya Foster, Lauren MacMullen, Jayamanna Wickramasinghe, Anastasia Lucas, Avi Srivastava, Steven Jacobson, Maureen E. Murphy, Samantha Soldan, Paul Lieberman, Noam Auslander

## Abstract

Human endogenous retroviruses (HERVs) are remmants of ancient infections which make up to ∼8% of the human genome. Their activity influences development, immunity, and cancer, but studying them has been limited by a key technical challenge: short-read sequencing cannot uniquely assign reads to these highly repetitive elements. Here, we present ERVmancer, a phylogeny-informed method that resolves the read-mapping ambiguity and quantifies HERV expression across scales, from individual loci to entire retroviral clades, depending on mapping confidence. Benchmarking with sample-matched long- and short-read data generated in this study demonsrates that ERVmancer outperforms existing approaches in both sensitivity and specificity. Application of ERVmancer recapitulates known HERV expression patterns in multiple sclerosis and uncovers new biology in breast cancer, including suppression of HERVH-LTR7 by p53. By enabling accurate and scalable quantification of integrated retroviral elements, ERVmancer provides a broadly applicable resource for investigating retroviral mechanisms in health and disease.

## Introduction

Accurate quantification of short-read RNA sequencing data is the first step for a thorough understanding of the relationship of expression and phenotype^1, 2^. For human protein-coding genes, this problem has been comprehensively addressed, with numerous tools for aligning short-read RNA reads to a specific gene locus allowing accurate quantification of these reads^3-6^. However, only about 2% of the human genome is annotated as protein-coding genes^7, 8^. The remainder of the genome is heavily enriched in highly repetitive sequences, including a class of repetitive sequence known as Human Endogenous Retroviruses (HERVs), derived from ancient retroviral infections^9-11^. Many of these HERV repetitive elements continue to express viral RNA^12-15^, and have been implicated in the progression of numerous diseases, including Multiple Sclerosis (MS)^16-20^, Systemic Lupus Erythematosus (SLE),^21-23^ other immune diseases^24-29^ and cancers^30-34^.

HERV expression has increasingly become a focus of interest for research to understand the role of HERVs in human health, cancer, immune regulation, and as prognostic biomarkers^15^. However, tools designed to align and quantify short-read RNA sequencing data for protein-coding genes fail to accurately map reads to the correct locus when high homology is present, severely limiting the use of traditional alignment tools and protocols for analysis of HERVs^35-37^. Previously, several tools have been developed using different approaches to quantify expression of HERVs and other repetitive elements of the genome, the most popular of which are TELEscope^37^ and ERVmap^36^. TELEscope is a general-purpose algorithm for quantifying repetitive regions, using Expectation-Maximization to assign ambiguous reads to specific loci. ERVmap uses a stringent alignment followed by a custom filtering approach to remove reads that share high homology with other HERVs. While these methods improve the mapping for highly expressed HERVs with unambiguous regions, many HERVs with fewer unambiguously mapped reads will be incorrectly assigned, especially those with low or moderate expression levels. In addition, integration events within protein-coding regions can lead to HERV sequence homology with non-target regions or protein coding genes, resulting in false positive HERV mapping with these methods^38, 39^. Other methods that have been developed, such as the Bayesian SalmonTE^40^, share similar issues to TELEscope in quantifying lowly expressed HERVs with high homology to non-target regions. In microbiology, lowest common ancestor (LCA) approaches based on phylogeny have been successfully used to assign ambiguous reads to taxa, leveraging the evolutionary conservation of each sequence.^41-44^

In this study, we present ERVmancer, a phylogenetic approach for resolving repetitive HERV reads in bulk RNA sequencing data, enabling accurate quantification of HERV expression and analysis of HERVs in human disease. We generated a benchmarking dataset by combining highly accurate and unambiguously mapped PacBio long-read RNA sequencing with sample-matched Illumina short-read RNA sequencing of lymphoblastoid cell lines from multiple sclerosis patients and healthy controls. Using this dataset, we show that ERVmancer outperforms existing methods in recapitulating HERV expression patterns from short-reads. Analysis of simulated data highlights misclassification patterns in prior approaches and demonstrates how ERVmancer resolves them. To illustrate its ability to uncover new biology, we applied ERVmancer to breast cancer cell lines, revealing a previously unrecognized role of p53 in suppressing a clade of HERVH-LTR7, which we experimentally validated. ERVmancer thus provides accurate, clade-level HERV quantification, enhancing biological interpretability and enabling a deeper mechanistic understanding of these complex repetitive elements in human disease.

## Results

### ERVmancer: leveraging phylogeny to quantify human endogenous retrovirus expression

To overcome the challenges of quantifying HERVs from short-read RNA sequencing data, we developed ERVmancer, a phylogenetic approach for HERV expression quantification. ERVmancer resolves issues of HERV homology by mapping ambiguously aligned short RNA sequencing reads onto a phylogenetic tree, assigning each read to the Lowest Common Ancestor (LCA) of its possible source HERVs (Figure 1).

**Figure 1.**
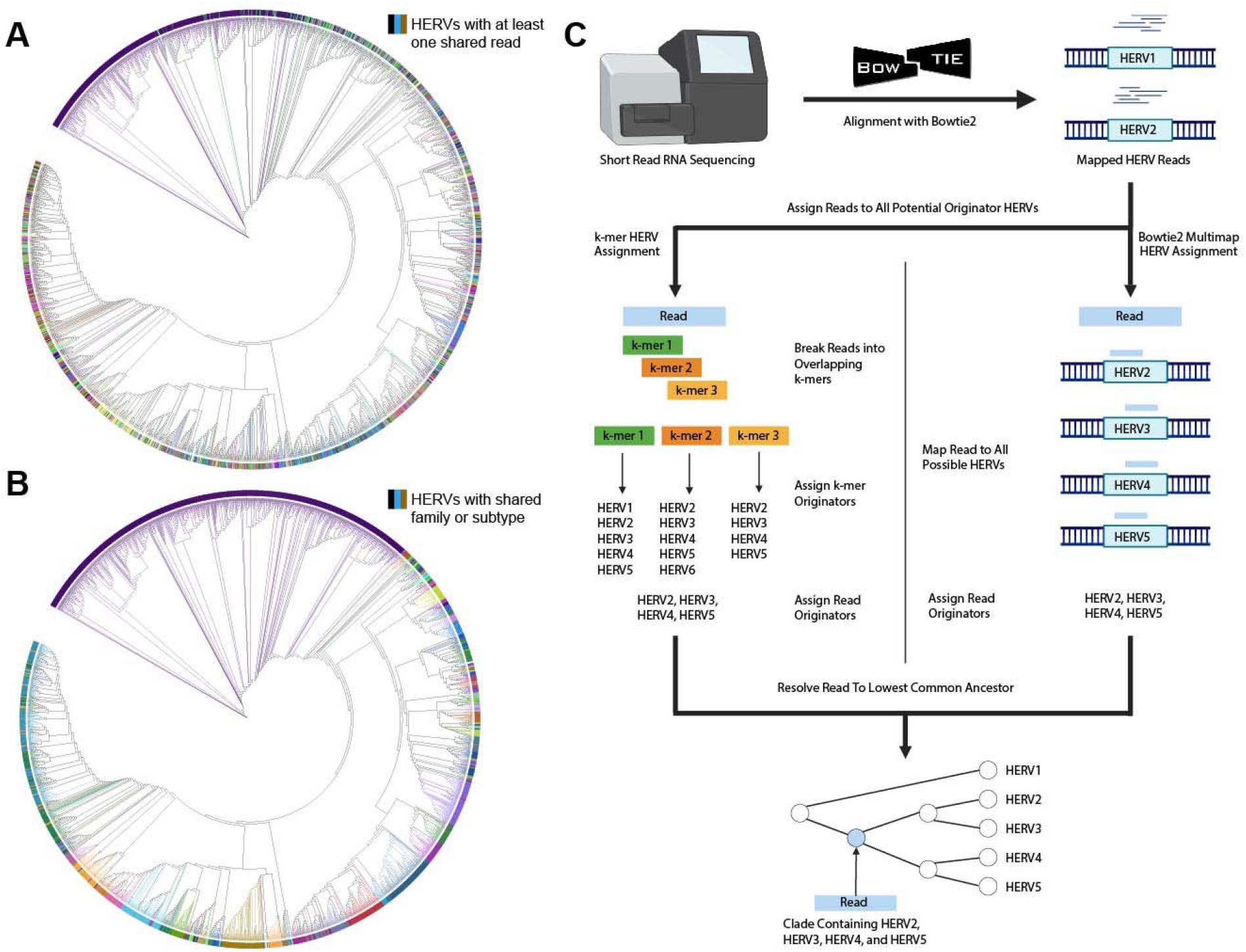
Overview of ERVmancer. **(A)** A phylogenetic tree of 3,043 potentially protein-coding HERVs. Branches are colored by the internal clade at which the cumulative mapping probability reaches 1. HERVs of the same color therefore share at least one identical read. **(B)** The phylogenetic tree colored according to the family or subtype identifier of the HERV. HERVs with the same family or subtype share the same color. **(C)** Workflow of ERVmancer assigning short RNA-seq reads to the HERV phylogenetic tree. Reads are first mapped to the hg38 genome, and those aligning to at least one potentially active HERV are retained. Each retained read is then assigned to a candidate set of originator HERVs using Bowtie2 multimapping and k-mer-based mapping and is placed in the phylogenetic tree at its LCA by combining the two strategies.

ERMancer’s underlying model consists of a phylogenetic tree constructed from 3,043 HERVs curated from the HERV database HERVd^45^. These HERVs were selected through a previously published filtering pipeline^46^ that requires the presence of at least one marker retroviral reading frame (Supplemental Table 1, Methods). The HERVs were used to infer a phylogenetic tree (Methods). The tree enables ambiguously mapped reads to be assigned to internal nodes, representing sets of possible source HERVs that share substantial sequence homology. To assess the effectiveness of this phylogenetic framework for read mapping, we generated a cumulative probability distribution capturing whether a read originating from a given HERV maps uniquely to that HERV or instead collapses into an internal node. This distribution reflects the degree of read overlap across HERVs in the tree (Methods). For a typical read length of 75 bp, we projected the cumulative probability distribution onto the phylogenetic tree, coloring the branches according to the point at which each HERV reached a cumulative probability of 1 (Figure 1A). Therefore, the colors indicate the worst-case resolution at which reads from each HERV can be distinguished. The CDF reflects the retroviral family structure and phylogeny, supporting the notion that different HERVs can be resolved at different scales depending on their sequence family and evolutionary relationships (Figure 1B).

Across the entire tree, HERVs achieve a cumulative probability of 1 at an average of 12.7 internal clades above the leaf node. The largest clade (purple in Figure 1A) illustrates both the difficulty of mapping short HERV reads and the advantage of our phylogenetic approach. This clade comprises 634 HERVs from the HERVH-LTR7 family, which on average share only 2.8% of their reads. Notably, this overlap does not correspond to a single large, evolutionarily conserved region. The most conserved 75-mer occurs in only 131 (20%) of these HERVs. Instead, this pattern arises from a combination of high sequence similarity and extensive evolutionary diversification within the family, which undermines purely alignment-based strategies (Supplemental Figure 1). By leveraging phylogeny, our method assigns each read to the lowest confidently resolved clade, preserving biological interpretability of multimapped reads at the clade level while minimizing both misclassification and loss of information.

In its default mode, ERVmancer processes raw FASTQ files from bulk short-read RNA sequencing (Figure 1B). Reads are first aligned to the hg38 reference genome using Bowtie2, and those mapping to at least one HERV locus are retained, filtering most of the non-target reads. Each retained read is then evaluated using two complementary strategies, a k-mer-based approach, and a multimapping-alignment approach with Bowtie2 across the HERV database (Methods, Supplemental Methods). In the k-mer approach, every read is decomposed into overlapping 31 bp k-mers, which are queried against the precomputed k-mer database. The set of possible originator HERVs for each k-mer is recorded, and the intersection across all k-mers from a read yields either a single HERV or a list of candidate originators. Because the k-mer database was prefiltered to remove sequences overlapping with non-target HERV k-mers, this step acts as an additional non-target filter. In the alignment approach, candidate originators are defined by all multimapping alignments reported by Bowtie2. Finally, ERVmancer integrates both strategies by assigning each read to the phylogenetic tree, to a leaf if only a single HERV was identified, or to the Lowest Common Ancestor (LCA) if multiple candidate originators remain.

### ERVmancer accurately recapitulates HERV expression patterns identified with long-read sequencing

To benchmark ERVmancer against existing methods, we generated paired short- and long-read (PacBio Isoseq) RNA sequencing data. The long-read data allows mapping HERV reads to individual loci, serving as a locus specific benchmark for evaluating short-read methods^47^. We sequenced 10 lymphoblastoid cell lines (LCLs) from individuals with MS and controls: three cell lines were from healthy controls (HC), three cell lines were from patients with active disease (AMS), and four cell lines were from patients from stable disease (SMS) (Methods, Supplemental Table 2). Given the longstanding implication of HERVs in MS pathogenesis, these samples provide a biologically relevant model with substantial variability in HERV expression for benchmarking purposes^17, 18, 48, 49^. For ground truth long-read mapping, full length non-concatemer (FLNC) reads were aligned to the human hg38 genome using minimap2 and reads mapping to the 3,043 HERV loci of interest were counted and normalized (Methods).

We benchmarked ERVmancer’s performance against two widely used methods for quantification of HERVs from short-read RNA sequencing: TELEscope^37^ and ERVmap^36^. All three methods were run on the same short-read MS cell line datasets, and all three methods were run using the GTF file of the 3043 HERVs to allow direct comparison between ERVmancer, ERVmap, and TELEscope. ERVmancer was run with default settings (Methods). To optimize performance for competing methods, the Bowtie2 step for TELEscope was run according to the parameters used in the HERV analysis section of the original TELEscope paper^37^ (Methods). All other TELEscope settings were set to default. ERVmap was run using default settings. To compare the counts yielded by the long-read data to ERVmancer’s phylogenetic output, we mapped the long-read counts to ERVmancer’s phylogenetic tree (Methods).

To assess how well different methods recapitulate HERV expression patterns captured by long-read sequencing, we calculated Spearman Correlation Coefficients between normalized counts per million (CPM) from short- and long-read datasets. Across all samples, ERVmancer consistently showed stronger correlations with long-read data compared to TELEscope and ERVmap (Figure 2A). A bootstrapping analysis involving repeated subsampling of 50% or 80% of the HERV features confirmed that these results were robust and not driven by outliers (Figure 2B,C). ERVmancer also outperformed TELEscope and ERVmap at the sample level, indicating that the improvements were not attributed to individual samples or phenotype differences (Figure 2D). This was further supported by direct comparison of minimap2 outputs against short-read methods, where ERVmancer again displays the strongest concordance with the long-read data (Figures 2E-G).

**Figure 2.**
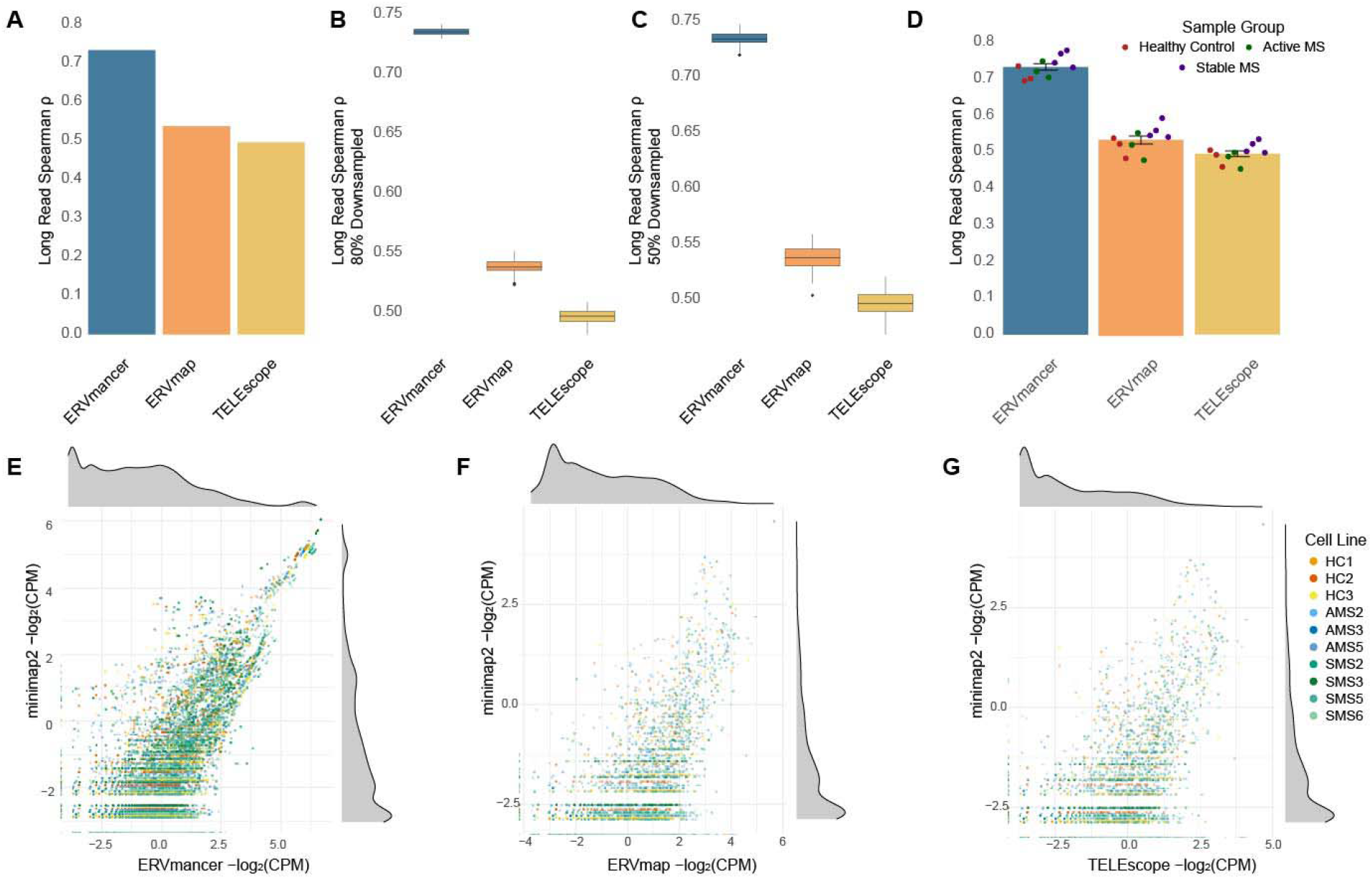
Benchmarking ERVmancer against previous short-read quantification methods using long-read data. **(A)** Spearman correlation coefficients comparing long-read HERV CPM with short-read quantification using ERVmancer (blue), ERVmap (orange), and TELEscope (yellow). **(B)-(C)** Spearman correlation coefficients from bootstrap analysis with random downsampling of HERV features 80% (B) or 50% (C). **(D)** Mean ± standard deviation of Spearman correlation coefficients between long-read CPM and short-read methods ERVmancer (blue), ERVmap (orange), and TELEscope (yellow). **(E)-(G)** Scatterplots of long-read based CPM (-log2 scaled) from minimap2 (y-axis) versus the short-read CPM (x-axis), ERVmancer (E), ERVmap (F), and TELEscope (G).

Collectively, these results demonstrate that ERVmancer more accurately recapitulates HERV expression patterns observed in long-read than existing quantification methods. As expected, ERVmancer also better captures phenotype-specific and HERV family expression patterns observed in long-read sequencing (Supplemental Figure 2, Supplemental Figure 3). Notably, across all methods, we observed a global increase of HERV expression in stable MS (SMS) relative to both active MS and healthy controls. The number of SMS-associated HERV clades and the limited cohort size precluded identification of individual HERVs suitable for validation. Despite this fact, we found evidence that SMS-activated HERVs were enriched on chromosome 19, near genes encoding KRAB-domain proteins, a family known to suppress herpesvirus infections and regulate viral latency^50, 51^. This consistent pattern suggests systemic HERV activation that may be sustained in B cells of SMS patients (Supplemental Figure 2).

**Figure 3.**
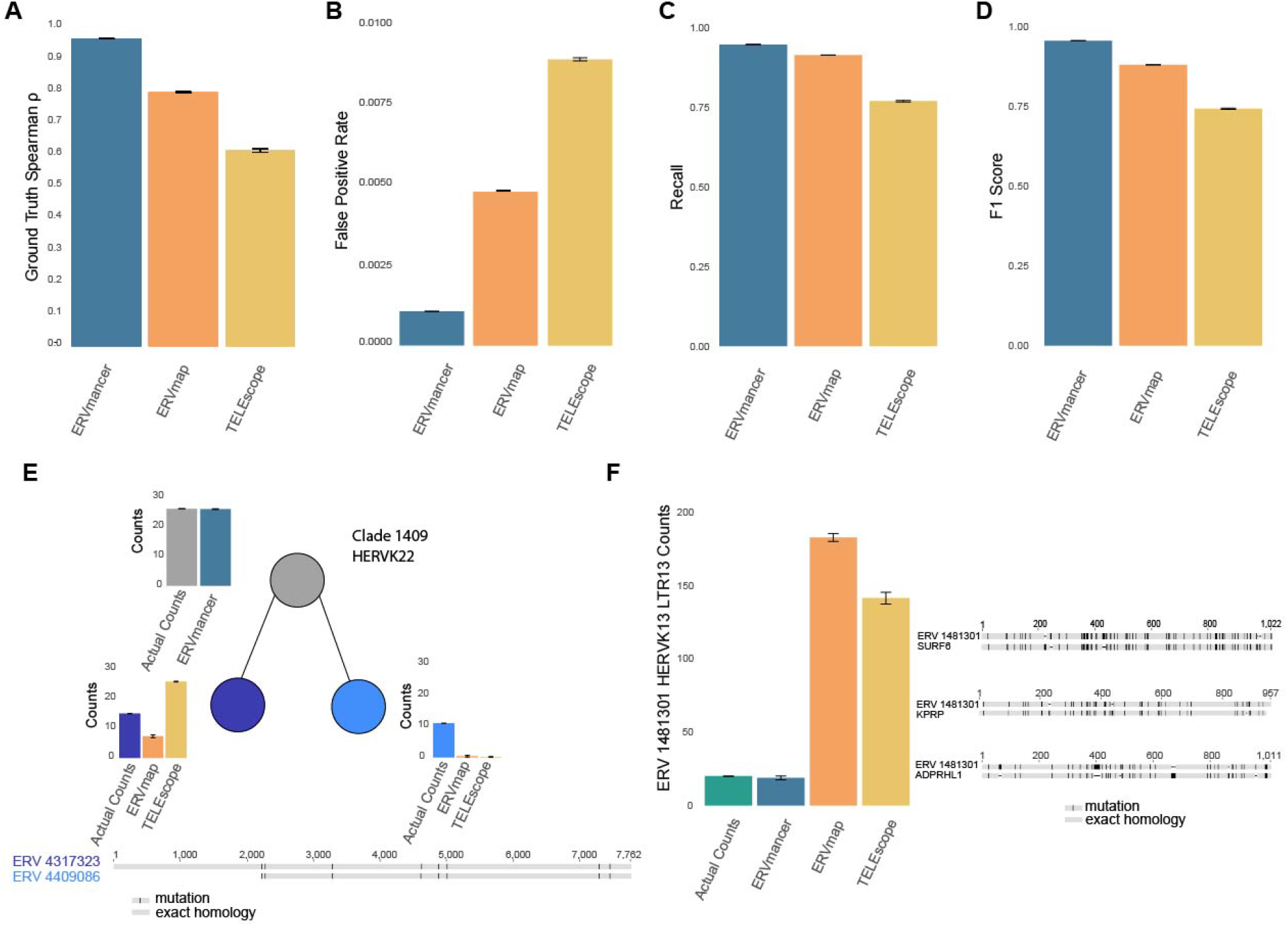
Benchmarking ERVmancer using simulated data. **(A)** Spearman correlation coefficients comparing ground-truth counts with short-read quantification by ERVmancer (blue), ERVmap (orange), and TELEscope (yellow). **(B)** False Positive Rate for each short-read method. **(C)** Recall for each short-read method. **(D)** F1 scores for each short-read method. **(E)** Comparison of simulated versus outputs from ERVmancer, ERVmap, and TELEscope, for clade 1409 containing two highly homologous HERVs, ERV 4317323 (chrX:148801576-148809340) and ERV 4409086 (chrX:148757259-148762800). **(F)** Comparison of simulated counts with ERVmancer, ERVmap, and TELEscope outputs, for ERV 1481301 (chr16:2710263-2720538), which shares homology with multiple protein-coding genes.

### Performance benchmarking using simulated short-read datasets

To further benchmark ERVmancer and assess its performance, we compared it to TELEscope and ERVmap on simulated data. We generated 10 datasets with one million reads each simulating the expression of the 3,043 potentially protein-coding HERVs with background transcriptomic reads from the gencode GRCh38 combined transcriptome^52^ using ART^53^. All methods were run under the same setup as in the real data benchmarking (Methods). ERVmancer showed consistently higher Spearman correlation with simulated reads across datasets (Figure 3A). It also achieved markedly lower false positive rate, less than a third of that from TELEscope and ERVmap, and higher recall (Figure 3B,C). Consequently, ERVmancer obtained the best F1 scores (Figure 3D). These results show that ERVmancer’s dual filtering steps more effectively remove non-target reads and improve HERV read mapping and quantification compared to existing methods.

Using the simulated data, we highlight examples where ERVmancer’s phylogenetic approach offers clear advantages. First, we examine a clade containing two highly homologous HERVs on chromosome X. ERVmap discarded reads mapping to these HERVs due to multimapping filters, while TELEscope incorrectly assigned nearly all reads to one of the HERVs (Figure 3E). In contrast, ERVmancer assigned all reads within the clade, yielding a correct count per clade. Second, we evaluated a case with high homology to non-target sequences, HERVK13 LTR13 on chromosome 16. TELEscope and ERVmap over-assigned reads by ∼6-8 fold compared to the simulated ground truth. ERVmancer’s k-mer filtering enabled accurate estimation of counts, despite considerable homology with protein-coding genes, such as *SURF6, KPRP*, and *ADPRHL1*, in addition to other HERVs and lncRNAs (Supplemental Figure 4).

**Figure 4.**
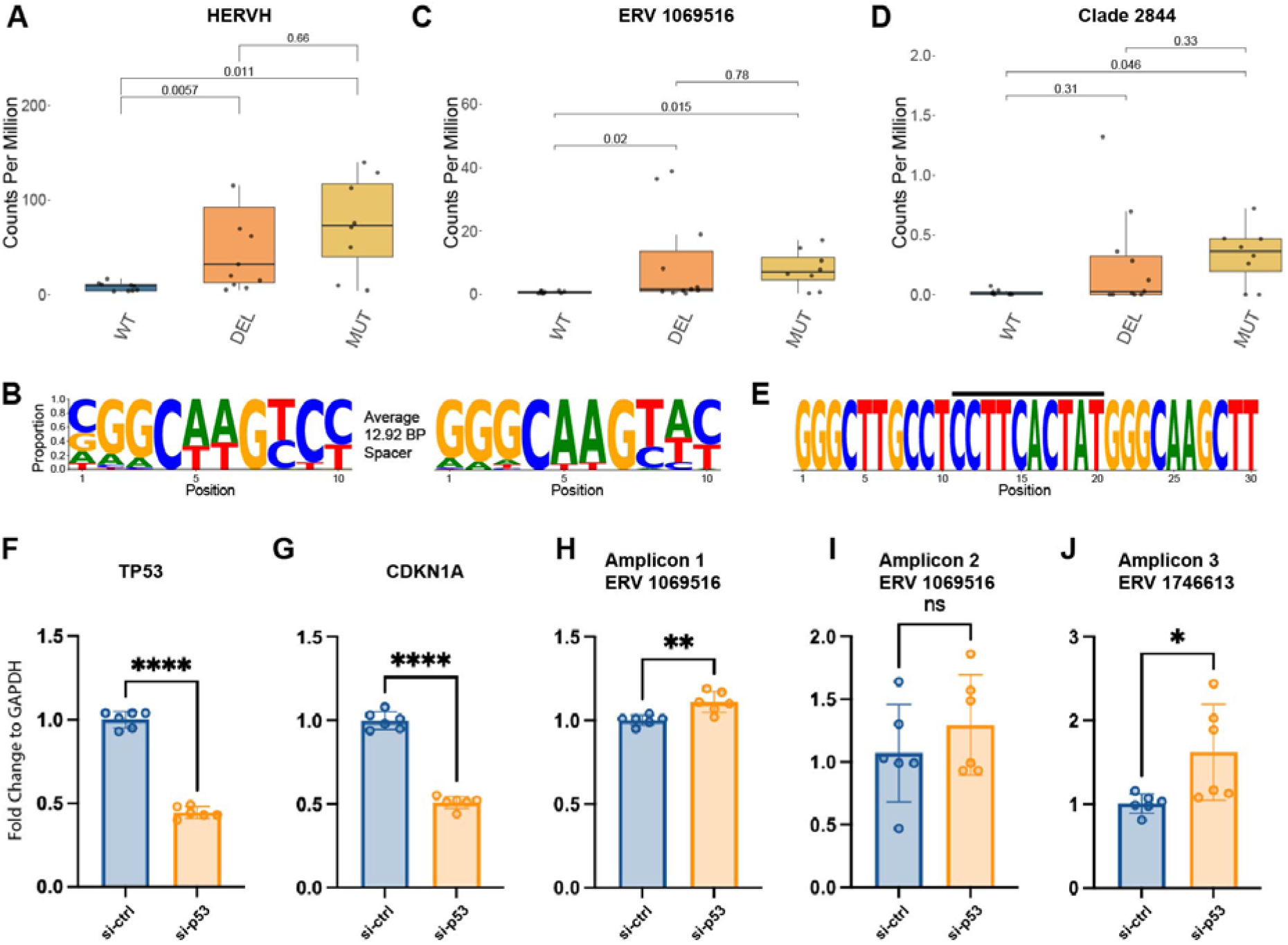
ERVmancer uncovers p53-mediated role in regulating HERVH-LTR7 expression in breast cancer cell lines. **(A)** Summed expression of HERVH clades across *TP53* states, showing higher levels in MUT and DEL compared to WT. MWU p-values are indicated. **(B)** Sequence logo representation of conserved p53 binding sites present in the 482 HERVH-LTR7s with p53 binding sites. **(C), (D)** Expression levels across p53 states for HERVH-LTR7 1069516 (chr13:109917438-109923506) (C) and internal clade 2844 (D), containing HERVH-LTR7 1746613 (chr18:70991850-70997625). MWU p-values across the states are indicated. **(E)** Sequence logo representation of conserved P53 binding site present in HERVH-LTR7 1069516 and 1746613. Black bar indicates the 10 bp spacer. **(F)-(J)** qPCR results using MCF7 cell lines with either control or silenced p53, *TP53* (F) *CDKN1A* (G) Amplicon 1 (H), targeting HERVH-LTR7 1069516 (chr13:109917438-109923506). Amplicon 2 (I), targeting HERVH-LTR7 1069516, and Amplicon 3 (J), targeting HERVH-LTR7 1746613 (chr18:70991850-70997625).

### ERVmancer unravels a new biological association of HERV clade with P53

To assess ERVmancer’s ability to capture biologically relevant HERV expression patterns, we applied it to RNA-seq from breast cancer cell lines in the Cancer Cell Line Encyclopedia (CCLE)^54-56^. Given prior evidence linking *TP53* to retroelement regulation^57-59^, we compared HERV expression across lines with wild type (WT) *TP53* (n=9), deleterious missense *TP53* mutations (MUT, n=8), or truncating/frameshift mutations (DEL, n=12).

We applied ERVmancer to the CCLE RNA-seq data and used a Mann-Whitney U (MWU) significance test to compare HERV clade expression across *TP53* states. Several HERVH-LTR7 clades showed consistent increase of expression in *TP53* MUT and DEL lines compared to WT, consistent with a suppressive role of p53 (Figure 4A). Binding-site analysis revealed conserved p53 binding motifs within these clades. Further, we found a significant enrichment between the HERVH-LTR7 types showing increased expression in *TP53* MUT and DEL lines, and those having canonical p53 binding sites (hypergeometric enrichment p-value = 5.23 x 10^-24^, Figure 4B, Supplemental Figure 5).

We next extracted reads from the HERVH-LTR7 clades with increased expression in *TP53* MUT/DEL and containing canonical p53 binding sites. From these, we assembled viral contigs and identified open reading frames (ORFs) matching GAG, POL, or ENV proteins, suggesting potential functionality. Two candidate HERVs emerged, HERVH-LTR7 ERV 1069516 (located on chr13) and HERVH-LTR7 ERV 1746613 (located on chr18) (Figure 4C-E). Notably, ERV 1069516 has been reported as a biomarker in lung cancer^60^ and osteoporosis^61^, while ERV 1746613 promotes radiotherapy resistance in breast cancer^62^. ERVmancer analysis showed both HERVs were suppressed in WT *TP53* and upregulated in MUT/DEL (Figure 4C-E). We validated these findings with three sets of 30 bp primers (Methods).

To test this experimentally, we silenced p53 using siRNA in the MCF7 cell line (Figure 4F, Methods). As expected, the level of the p53 target gene CDKN1A decreased with p53 siRNA, confirming effective p53 knockdown (Figure 4G). Conversely, the expression of ERV 1069516 showed modest increase, and ERV 1746613 showed more considerable increase, after p53 silencing (Figure 4H-J). These results validate ERVmancer’s prediction of a suppressive role of p53 in regulating HERVH-LTR7 expression and highlight ERVmancer’s ability to uncover novel expression patterns in evolutionarily conserved HERVs relevant to human diseases.

## Discussion

Quantification of repeat regions using short-read RNA sequencing is challenging but critical for understanding their role in human diseases. HERVs are a key class of repeats that can express proteins and modulate immunity and patient outcomes. Leveraging both existing and newly generated short-read RNA sequencing data provides a rich resource for studying HERVs and their effects on multiple disease states. To link HERV expression to human conditions, we developed ERVmancer, a new approach that quantifies HERV expression by mapping short reads to a HERV phylogenetic tree. Using matched short- and long-read RNA-seq of LCLs from MS patients and controls, we show that ERVmancer better recapitulates HERV expression patterns than existing methods. On simulated data, ERVmancer also reduces false positives and false negatives, due to the k-mer filtering and phylogenetic mapping of highly homologous HERVs. Finally, we demonstrate its ability to unravel new biology by identifying and validating a role for p53 in the negative regulation of HERVH-LTR7.

We explored other approaches to quantify HERV expression, including clustering- and consensus-based methods (Supplemental Methods). However, variability in sequence heterogeneity and mutation patterns across HERV families prevented biologically meaningful grouping. Instead, we adopted a phylogenetic approach, which provides a flexible resolution to map both unique and highly homologous reads while preserving biological interpretability.

ERVmancer outperforms TELEscope and ERVmap in capturing HERV expression patterns in both long-read and simulated data. TELEscope’s use of local alignment with higher mismatch tolerance may increase its false positives in a way that the EM step cannot fully correct, suggesting that parameter search may benefit HERV analysis. ERVmap, lacking a genomic filtering step^63^, also produces more false positives than ERVmancer. While ERVmancer compares favorably, it is designed as an alternative for quantifying phylogenetic groups of protein-coding HERVs, rather than a full replacement for existing methods.

Limitations of our approach include the fixed phylogenetic tree, which is predefined and restricts quantification to the 3,043 HERVs with protein-coding potential. While these HERVs are particularly relevant to human disease, future work could expand the tree and k-mer dictionary to include additional elements. Another drawback is that differential expression analysis on a phylogenetic tree violates the independence assumptions of standard methods like DESeq2^64^, since nodes depend on their children. Development of phylogeny-aware differential expression methods tailored to HERV trees is needed to enable accurate clade-specific analysis^65-70^.

In conclusion, we developed ERVmancer, a phylogenetic method for mapping short-read HERV RNA sequencing. We show that it outperforms current standard methods and can uncover new roles of conserved HERV clades. While long-read seq can offer more definitive HERV locus assignment, the continued demand to enhance exisiting, more accurate and affordable short-read RNA sequencing underscores the value of ERVmancer for improved read assignment. In the future, ERVmancer may be adapted to enhance HERV read mapping across various data types, substantially improving our ability to investigate HERV molecular mechanisms.

## Methods

### Establishing a set of HERVs with protein-coding potential

HERVs with intact frames encoding POL/RT, GAG, or ENV were previously extracted and used here^46^. This criteria was used to improve phylogenetic influence and to select HERVs with potentially produce protein coding genes which could interact with the immune system. Briefly, genomic locations of reported HERV elements were downloaded from the HERVd HERV annotation database (https://herv.img.cas.cz)^71^, and the nucleotide sequences in hg19 were extracted using twoBitToFa^72^ and lifted into hg38. For each coding frame, blastx was employed against NR with an E-value cutoff of 1E-4, as well as a profile search^73^ against collected POL proteins annotated in GenBank in lentiviruses (as of September 2016) and aligning their amino acid sequences using MAFFT^74^. Sequences with at least one identified retroviral protein motif were extracted, yielding 3,043 HERVs that were used for our reference database. HERVs in this article are reported using their hg19 locations.

### Constructing a phylogenetic tree of HERVs

We performed a multiple sequence alignment (MSA) where the 3,043 HERVs were aligned together using MAFFT^74, 75^ alignment software v7.490. Using the MSA, a phylogenetic tree of all HERVs was established by employing FastTree^76, 77^ v2.1.11 with default parameters. The resulting unrooted tree was processed in python using Biopython^78^ v1.84 Phylo module and networkx^79^ v3.2.1. To increase the speed of ERVmancer, the tree was converted into two hash tables. The first hash table contains each clade as a key and all sub-children of that clade as values, and the second contains the leaves of the tree as keys and the clades back to the highest clade as values. The first hash table allows for quick summation of all the counts within a clade and all subchildren of a clade. The second hash table allows a list of HERVs to be rapidly assigned to the lowest clade containing all HERVs in the list.

### Mathematical framework for phylogenetic read assignment and resolution

To mathematically illustrate how the phylogenetic tree enables optimal resolution of all potential HERV reads, we defined a probability mass function describing the likelihood that a read from a HERV is mapped either to that HERV or to one of its ancestor clades. For HERV *h* in the reference database, let *r* be a read of length *k* from the set of all overlapping reads within *h*. The total number of such reads for *h* is *lengh*(*h*) – *K*+1.Denote the leaf node assigned to h as *n*_0_, and let *n*_*i*_represent the clade corresponding to the *ith* ancestor of *n*_0_.

A read *r* is assigned to *n*_0_ if it is a unique substring of *h* across the leaves of the HERV phylogenetic tree. If *r* is not uniquely found in *h*, i.e., *r* is a substring of additional HERVs, we iteratively check each higher clade *n*_*1*_ through *n*_*i*_, assigning *r* to *n*_*i*_ if *r* is unique within all HERVs contained under *n*_*i*_, meaning *r* does not appear in any HERV outside of *n*_i_ or its descendants. Each read *r* is uniquely assigned to one node in the tree. We thus define each HERV-clade pair (*h,n*_*i*_)as *x*.

The probability mass function of the HERV-clade pair *x*=(*h,n*_*i*_)is defined as

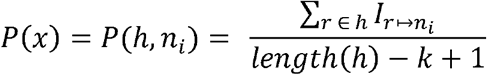

Where *I*_*r*↦*ni*_ is the number of reads *r* uniquely assigned to clade *n*_*i*_. Thus, the probability mass function defines the probability any read from *h* is directly assigned to *n*_*i*_.

The cumulative distribution for pair *h,n*_*t*_ is defined over all pairs *h,n*_*t*_ up to clade *n* (*t* ≤*i)* is:

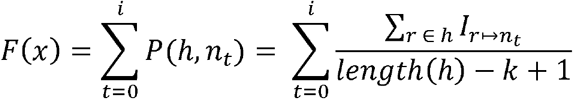

Because every read *r* from HERV *h* is assigned either to *h* itself or to an ancestor clade in the tree, the total probability sums to 1.

To evaluate the likelihood of the worst possible case where a HERV is assigned to the highest possible level in the phylogenetic tree, we calculated the cumulative distribution when read size *k* = 75 (corresponding to the worst case for single-end sequencing with read length of 75 bp). For each HERV, we calculated the node at which the HERV’s cumulative distribution first reaches the upper bound of 1, representing the level of the tree to which any read of that HERV would be mapped. We then colored all HERVs sharing the same lowest cumulative distribution upper bound clade. Therefore, HERV leaves with unique colors represent HERVs with no shared reads in the tree, while HERVs with shared colors represent a clade with potential reads multimapping to HERVs in that clade. These colors were then plotted onto the phylogenetic tree using ITOL v6^80^.

### ERVmancer alignment step

ERVmancer in its default mode receives as input single- or paired-end RNA-seq FASTQ files. These FASTQ files are aligned using Bowtie2 with paired end sequencing parameters:

--very-sensitive --end-to-end -X 1500 --no-mixed --no-discordant --no-dovetail --no-unal --score-min L,-0.1,-0.1 -k 100

And single end sequencing parameters:

-N 1 -L 10 --very-sensitive --end-to-end --no-unal --score-min L,-0.1,-0.1 -k 100

These parameters are used as default within the ERVmancer package. An additional entry point of ERVmancer allows end users to provide their own Bowtie2 SAM file output as input for ERVmancer to allow running with user-defined Bowtie2 parameters (see GitHub package documentation). In either entry point, the SAM file is converted to a BAM file using samtools^81^ v1.20 and then subsetted to include only potentially protein-coding HERV regions with bedtools. These HERV regions are defined using the hg38 locations of the 3,043 HERVs of interest from HERVd^45, 71^ and is available on Zenodo through the ERVmancer GitHub.

### ERVmancer k-mer-based read mapping

To map HERV reads using k-mers, we constructed a hash table of every possible 31 base pair overlapping k-mers and their reverse complements found in the 3,043 likely protein-coding HERVs. The choice of k = 31 was made to reduce the likelihood of error k-mers while increasing the number of k-mers per read, consistent with prior studies^41, 82, 83^. Concurrently, we prepared another 31 bp k-mer hash table from the gencode hg38 transcriptome sequences (https://www.gencodegenes.org/human/^52^). Because the gencode hg38 transcriptome contains HERV transcripts, we removed any transcripts which shared 60% of the k-mers present in our HERV k-mer database. Next, we removed any k-mers from the HERV k-mer hash table which appeared in the filtered hg38 transcriptome hash table. This filters out potential non-target false positive k-mers from the HERV k-mer database. We created the final hash table by assigning each k-mer hash to a list of HERVs that could have produced the k-mer.

Through the read assignment, reads which Bowtie2 maps to HERV regions are assigned to a list of potential HERVs using the k-mer hash table. Each read is broken down into all overlapping 31 bp k-mers, and every k-mer in the read is used as a hash to return a HERV or list of HERVs from which the k-mer could have originated. After all k-mers are mapped, the intersection of the HERV lists is taken, allowing more unique k-mers to contribute more information for mapping the read as a whole. For reads whose k-mers yield an empty intersection, the union of HERV maps are returned to address mutations that may affect some of the k-mers. Reads which do not have any k-mers in the hash table are removed from consideration entirely to discard reads which are most likely not from the target HERVs. Reads which have any k-mers present in the HERV hash table are retained, accommodating a moderate number of mutations. The HERV or set of HERVs identified from the read’s k-mers are then mapped to the phylogenetic tree. If the read maps to a single HERV, it is assigned to the corresponding leaf. If the read is mapped to multiple HERVs, it is assigned to the lowest internal clade that encompasses all of them.

### Bowtie2 output phylogenetic mapping

Concurrently, we directly use the Bowtie2 multimapped read assignments to map reads to the phylogenetic tree based on the alignment output. For each read, all HERVs assigned by Bowtie2 are collected into a list. Similarly to the mapping employed by the k-mer step, if the read maps to a single HERV, it is assigned to the corresponding leaf, whereas a read mapped to multiple HERVs is assigned to the lowest internal clade that encompasses all the originator HERVs.

### Incorporating the alignment and k-mer outputs to yield tree counts and normalized counts

The final output uses a combined approach between the k-mer and multimapped phylogenetic assignments. We observed that using the Bowtie2 assignments generally results in lower LCAs than the k-mer approach; however, the Bowtie2 assignments also result in higher false positives, as this step lacks non-target filtering (Supplemental Figure 4A,B). To achieve the most specific assignment of reads into HERV phylogeny while reducing false positives, the final tree assignments use the lowest assignment per read between the k-mer and Bowtie2 assignment methods, removing reads which did not have any HERV specific k-mers. The reads are normalized to the total counts of the original R1 FASTQ file, to establish CPM for each HERV clade. Finally, four outputs quantifying expression across the HERV trees are provided as output: (1) Raw counts of reads which were directly assigned to each node in the HERV tree, (2) The subtree sums of the raw counts for every node in the tree, representing the sum of reads assigned to any node and all of its descendants. Outputs (3) and (4) are the normalized CPM versions of outputs (1) and (2), respectively.

### Long-read and short-read preparation

Spontaneous lymphoblastoid cell lines (SLCLs) were generated ex vivo from MS patients and healthy controls (HC) as described previously^84^ (and PMID: 37562974). Importantly, SLCLs are distinct from traditional LCLs because they are transformed by the individual’s endogenous EBV, rather than using lab strain virus (e.g. B95.8). For this study, a total of ten SLCL lines were characterized; these lines were obtained from three healthy blood donors (HC1-3), three MS patients with active disease as determined by magnetic resonance imaging (AMS 2,3, and 5), and four patients with stable disease (SMS 2, 3, 5, and 6) (Supplemental Table 2). Notably, two pairs of cell lines were generated from the same individuals during different phases of their disease (AMS2= SMS6 and AMS5=SMS2). This cohort was used as a pseudo-gold standard to benchmark method’s ability to detect and quantify patterns of HERV expression within these cells. SLCLs were maintained in growth medium [RPMI-1640 medium (Gibco, Gaithersburg, MD) with 10% fetal bovine serum (FBS; Gibco, Gaithersburg, MD), 1% gentamicin (50mg/ml; Quality Biological, Gaithersburg, MD), and 1% L-Glutamine (200mM; Quality Biological)]. Total RNA was isolated from 2□× □10^6^ cells using the RNeasy Mini Kit (Qiagen, Germany) and treated with deoxyribonuclease (Qiagen, Hilden, Germany), following the manufacturer’s protocol. An RNA integrity number (RIN) value greater than 9.0 was determined by TapeStation (Agilent Technologies) for all samples prior to short paired-end and long-read RNA-seq. For paired-end RNA-seq, sequencing library preparation for SLCLs was completed using the True-seq library preparation kit (Illumina, San Diego). Sequencing was performed with an Illumina NextSeq 500 system in high-output mode to generate ∼15-20□× □10^6^ reads (2□× □150□bp) across three multiplexed and pooled samples. For long-read RNA-seq isoform libraries were prepared with the PacBio Iso-Seq Express kit according to the manufacturer’s protocol, which included oligo(dT)-primed cDNA synthesis and SMRTbell adapter ligation. Barcoded libraries were pooled equimolarly and sequenced on the PacBio Revio platform (Mount Sinai, NY), targeting ∼4–5 million HiFi reads per sample.

### Mapping PacBio long-read sequencing data

To obtain the pseudo-gold standard of long-read HERV expression mapping for benchmarking methods that map short-read sequencing, we aligned the FLNC BAM files from PacBio with minimap2^85^ v2.28 using the hg38 reference genome from UCSC^72^, with the following command:

minimap2 -ax splice:hq -B 4 --sam-hit-only -uf

All other parameters set to default. The SAM file output was then filtered for only high-quality alignments using the command:

samtools view -F 4 -q 30 -F 256

We used FeatureCounts^86^ (subread v2.0.6) and HERVd genomic locations to obtain count data of the long-reads. We additionally overlaid the counts onto the phylogenetic tree by mapping counts to the HERV leaves and summing up the phylogenetic tree to yield clade-level counts. CPM was calculated using the original FLNC BAM file.

To quantify HERV expression from short-reads for the benchmarking comparisons, we ran ERVmancer with default parameters on the short-read RNA sequencing FASTQ files for every sample. HERV count data was normalized into CPM, as described above.

### Employing TELEscope and ERVmap for benchmarking comparisons

TELEscope^37^ was installed through conda according to the package instructions. Following the original paper’s specifications for HERV analysis, Bowtie2 was used for the initial alignment, with parameters as defined in the TELEscope paper for HERV quantification:

--very-sensitive-local -X 1500 --no-mixed --no-discordant --no-dovetail --no-unal --score-min L,0,1.6 -k 100.

All other Bowtie2 parameters set to default. TELEscope itself was run using default parameters according to the documentation. We provided the GTF file containing the hg38 locations of the 3,043 HERVs of interest from HERVd. Output was normalized to CPM using the total counts of the original R1 FASTQ files for each sample.

ERVmap^36^ was downloaded from GitHub (https://github.com/mtokuyama/ERVmap). We downloaded all dependencies according to ERVmap’s GitHub. We modified the ERVmap auto script to intake the GTF file for the 3,043 HERVs of interest in this study, otherwise ERVmap was run according to the instructions on GitHub. The output of ERVmap was normalized to CPM using the total counts in the original R1 FASTQ files for each sample.

To compare the performance across methods we calculated Spearman correlation between the long-read and short-read output using the spearmanr function from the scipy^87^ package (v1.13.0) in Python. For correlation between minimap2 output and ERVmancer, we used the full normalized summed count tree for both minimap2 and ERVmancer and flattened all internal clades and leaves into a vector which was used for the correlation calculation. For the correlation calculation between minimap2 and the other short-read methods, we used a single vector which contained read counts per HERV. For the bootstrapping analysis, we downsampled by selecting 50% or 80% of the HERVs or internal clades from the phylogenetic tree, then removed those HERVs or clades from consideration between minimap2 and ERVmancer. We only removed the HERVs from consideration between minimap2 and TELEscope and minimap2 and ERVmap, as these two methods do not use the phylogenetic tree. We randomly downsampled 100 times each for the 50% and 80% total HERVs selection criteria.

### Generating simulated data for benchmarking comparisons

10 simulated datasets of ∼ 1 million reads each were generated using the simulation tool ART with commands:

art_illumina -ss HS25 -l 150 -f 0.6 -m 200 -s 10 -na -rs (random seed per dataset)

All other commands were set to default. The input file used to generate the simulated datasets was a combined fasta of gencode GRCh38 transcripts with exact matches to the 3,043 HERVs removed and a fasta of the 3,043 HERVs of interest from HERVd.

### Simulated data analysis

ERVmancer, TELEscope, and ERVmap were run as above. ERVmancer was configured to return the exact assignment per read in addition to the count output, allowing direct comparison between the assignment and the actual read source. Due to the differences in read assignment between the three methods, we define True Positives (TP), False Positives (FP), and False Negatives (FN) differently for ERVmancer as compared to TELEscope and ERVmap. An ERVmancer read assignment is considered a TP if the read was assigned directly to the source HERV, or if the internal clade assignment contains the source HERV. An ERVmancer read assignment is considered a FP if the read was not assigned to the HERV leaf or to an internal clade containing the HERV but was assigned to a different HERV or a clade that does not contain the correct HERV. FN are defined as a HERV read which was not mapped to a HERV or HERV clade.

For TELEscope and ERVmap, we implemented a less stringent read assignment for evaluation. FP are defined as any assigned counts higher than the simulated HERV counts, and FN are defined as any assigned counts lower than the simulated HERV counts. TP are any assigned counts exceeding the simulated count values.

### CCLE cell lines analysis

We downloaded RNA sequencing data for 28 breast cancer cell lines from the CCLE database (Supplemental Table 3) using the Broad DepMap portal^56^ (https://depmap.org/portal) and the SRA toolkit^88^. Nine samples had an unmutated wild type p53, eight samples had a p53 mutation, and twelve contained a frameshift or premature stop codon in p53.

We compared RNA expression between the different conditions using an MWU test (scipy.stats.mannwhitneyu v1.13.0). We extracted out the reads which were assigned to HERVs or clades with unadjusted significant p-values between the mutant and WT p53 states.

We identified p53 binding sites using a custom python program which allowed searching for up to one mismatch in the first three or last three bp in the two palindromic flanking regions, and a variable 0-20 bp spacer in between the two palindromic flanking regions. Enrichment of HERVH-LTR7 loci with p53 binding sites was performed with a hypergeometric enrichment test (scipy.stats.hypergeom.sf v1.13.0).

### Primer development for experimental validation of p53 association

First, we simulated a potential primer experiment to test potential changes in expression of these reads by reconstructing the expected differentially expressed reads using SPAdes^89^ v3.15.5. The assembled constructs were converted into a Bowtie2 index and the original FASTQ files were realigned to this index. Potential differential expression was then reconfirmed using an MWU test as above, and the contigs with potential changes in expression were reconstructed again using SPAdes. The contigs were manually examined using ExPASy Translate^90^ and HHPred^91^ for the presence of an open reading frame of a HERV protein. We selected two candidate viruses that met these criteria, HERVH-LTR7 ERV 1069516 (chr13:109917438-109923506) and HERVH-LTR7 ERV 1746613 (chr18:70991850-70997625)..

We prepared 30 bp primers for these two candidate viruses (Supplemental Table 4). To generate unique primers from these regions with high homology to other areas of the genome, we identified 30 bp regions of overlap between the original SPAdes constructed contig and the second SPAdes constructed contig. We then blasted each of these 30 bp regions and only retained regions which were unique to the virus of interest. From these, we selected forward and reverse primers that had a melting temperature within 3°C as calculated using the New England Biolabs Tm Calculator (https://tmcalculator.neb.com/#!/main). We created three sets of primers: two for ERV 1069516, and one for ERV 1746613.

### Experimental validation of the predicted role of p53 in inhibiting HERV-LTR7

1,000,000 MCF7 cells were plated per condition (si-ctrl & si-p53). The next day, si-ctrl and si-p53 was added using Lipofectamine RNAiMAX Transfection reagent following the manufacture’s protocol. Following 24 hours of si-RNA treatment, cells were harvested, and RNA was isolated using the RNeasy Qiagen kit. RNA was then reverse transcribed using the High-Capacity Reverse Transcription Kit using the manufacture’s protocol. qPCR was performed on biological replicates with three technical replicates using the QuantStudio 5 machine and PowerUp SYBR Green Master Mix. Expression levels of the amplicons are expressed as fold change normalized to GAPDH. Materials used: RNeasy Kit (Qiagen – Cat#74106), PowerUP SYBR Green Master Mix (Thermo Fisher Scientific – Cat#A25742), Lipofectamine RNAiMax Transfection Reagent (Thermo Fisher Scientific – Cat# 13778150), High-Capacity cDNA Reverse Transcription Kit (Thermo Fisher Scientific – Cat# 4368814).

## Data and Code Availability

ERVmancer package and source code are freely and publicly available on the GitHub (https://github.com/AuslanderLab/ERVmancer), with detailed download and installation instructions on Bioconda. Scripts to reproduce key analysis and figures are also provided.

## Contributions

N.A. initiated the study. A.P. and N.A. performed research, analyzed the data, created the method, and wrote the manuscript. B.D. and A.P. created the ERVmancer Bioconda package. M.F., L.M., and M.M. performed cell culture and experimental validation. L.Y., S.S., S.J, and P.L. prepared long- and short-read RNA sequencing data on the MS samples. AL assisted in manuscript writing and mathematical notation. A.S. advised on methodology. J.W. assisted in code creation and technical support.

## Acknowledgments

We thank McKenna Reale and Timothy Kossenkov for assistance in testing the ERVmancer Bioconda package. Funding for this study was provided by the W. W. Smith Charitable Trust number C2308, the Michelson Medical Research foundation, and the R01 LM014503-02 from NIH/NLM. A.P. was supported by Wistar’s T32 grant 5T32CA009171.

