## Supplementary Methods and Figures for "A phylogeny-guided framework for decoding mechanisms of human endogenous retrovirus regulation in health and disease"

### Supplemental Table Legends

**Supplemental Table 1.** Metadata on the 3043 HERVs used by ERVmancer with at least one intact GAG, Pol/RT, or ENV protein, their hg19 genomic locations, and the closest gene to the HERV. HERV names are from HERVd.

**Supplemental Table 2.** Metadata on the spontaneous lymphoblastoid cell lines from Multiple Sclerosis patients and healthy controls.

**Supplemental Table 3.** Metadata on the CCLE breast cancer cell lines.

**Supplemental Table 4.** Metadata for primers used in figure 4.

### Supplemental Methods

#### The parallel ERVmancer assignment steps

ERVmancer uses two parallel methods to assign originator HERVs: a step using hashed k-mer tables, and a step using Bowtie2 multimapping assignments. These parallel approaches result in a more accurate tree assignment. After the initial Bowtie2 HERV read selection, the k-mer step removes more false positives (Supplemental Figure 4A) but tends to assign reads further from the leaves compared to the multimapping approach (Supplemental Figure 4B). This pattern is due to the k-mer step relying on the smaller, less informative 31 bp k-mers for assignment. In contrast, Bowtie2, by considering information across the entire read, tends to assign reads lower down in the tree (Supplemental Figure 4B). However, the multimapping step does not filter false positives, as it does not consider if the read also aligns to non-HERV regions. Therefore, the final read counts are made by removing all reads which are filtered in the k-mer step, and then assigning the read to the tree to the lowest LCA assignment between the k-mer and Bowtie2 multimapping approaches. The result is the best resolution of a HERV read to its LCA (Supplemental Figure 1A,B).

#### Examining alternative strategies for HERV quantification

Our initial attempts to solve the issue of quantification of HERV expression used clustering and consensus approaches. First, we applied CD-Hit-EST[1] to cluster reads from the 3,043 annotated HERVs and grouped these clusters as potential biomarkers. However, the resulting clusters lacked the necessary granularity for effective biomarker development or discovery, and their high heterogeneity prevented the construction of biologically meaningful consensus sequences.

Next, we explored a consensus-based alignment-informed approach using MAFFT[2, 3] to align every possible HERV pair, yielding ~4.7 million pairwise alignments. While homology was frequently observed between individual pairs, it did not extend consistently across groups of related HERVs, again precluding the derivation of accurate consensus sequences or coherent clusters.

Given these limitations, we turned to a phylogenetic approach. This framework offers a sliding scale of resolution, uniquely mapping reads can be directly assigned to specific HERVs, while reads from highly homologous regions can be placed at the most informative internal nodes. In doing so, the phylogenetic method preserves biological interpretability within the context of the inherent uncertainty of highly homologous reads.

**Supplemental Figures**

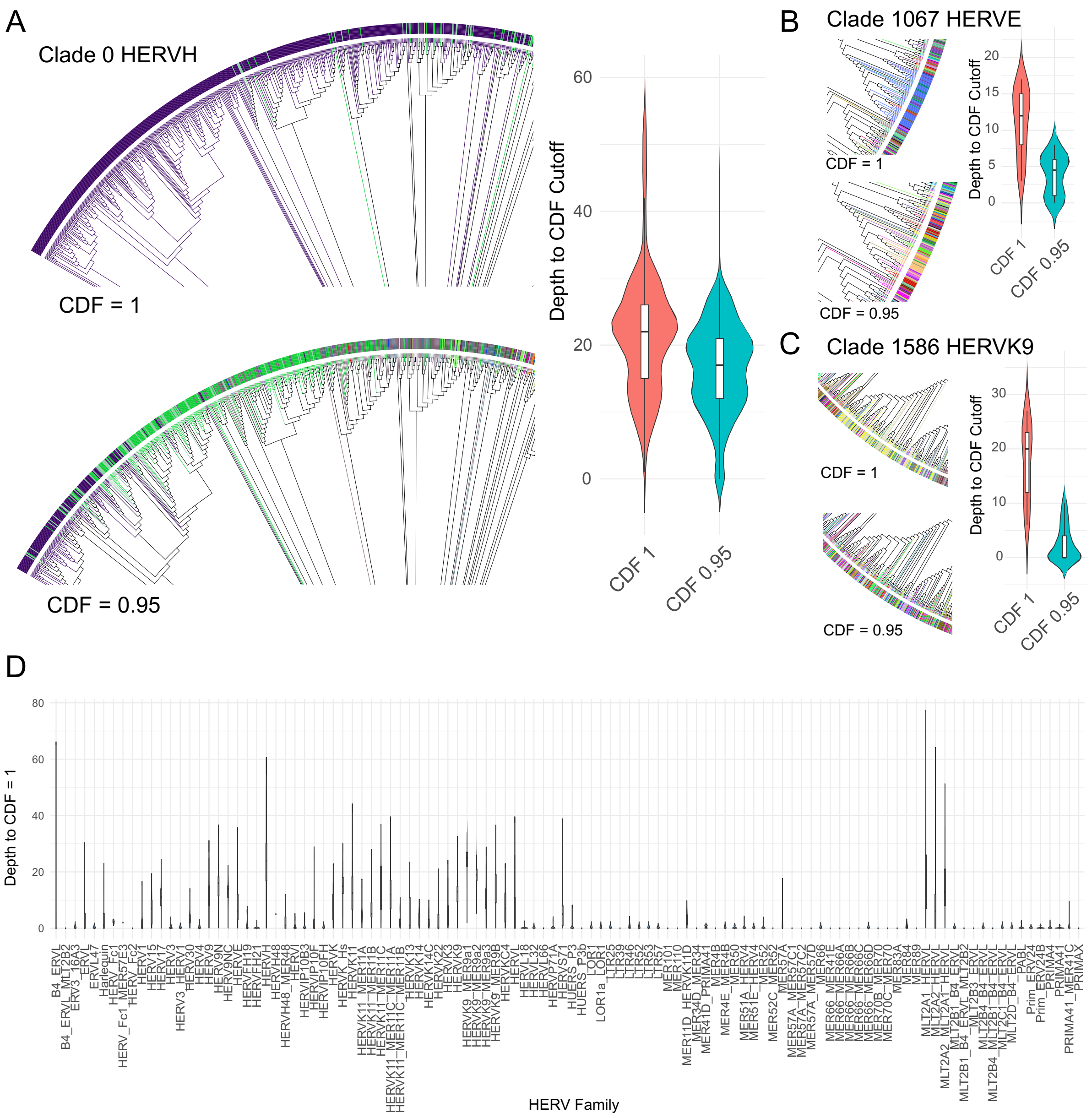


**Supplemental Figure 1. CDF specific patterns across HERV families and clades. (A)** Left: Slice of the tree comparing HERVH colored by a CDF cutoff of 1 vs a CDF cutoff of 0.95. Right: The depth from the HERV leaf to the specified CDF cutoff for HERVH.**(B)** Left: Comparison of Clade 1067 containing HERVE colored at CDF cutoff of 1 and CDF cutoff of 0.95. Right: Depth to reach the specified CDF forHERVE. **(C)** Left: Comparison of Clade 1586 containing HERK9 at CDF cutoff of 1. Right: Depth to reach the specified CDF for HERVK9 **(D)** Violin plots of the depth each HERV achieves a CDF of 1. HERVs are grouped by family or subtype.

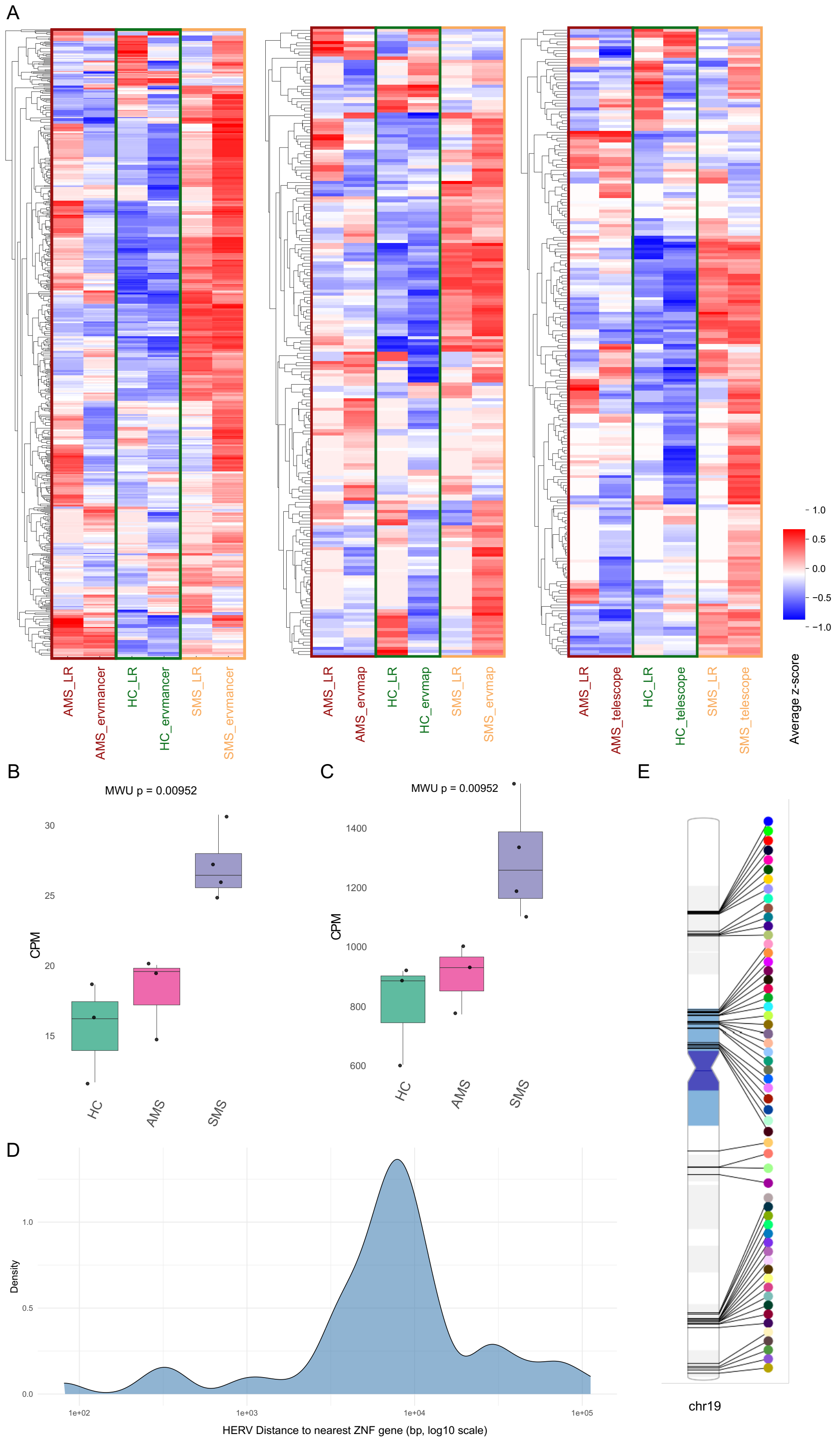
A
AMS_LR
AMS_ervmancer
HC_LR
HC_ervmancer
SMS_LR
SMS_ervmancer
AMS_LR
AMS_ervmap
HC_LR
HC_ervmap
SMS_LR
SMS_ervmap
AMS_LR
AMS_telescope
HC_LR
HC_telescope
SMS_LR
SMS_telescope
1.0
0.5
0.0
−0.5
−1.0
Average z-score
B
MWU p = 0.00952
CPM
30
25
20
15
HC
AMS
SMS
C
MWU p = 0.00952
CPM
1400
1200
1000
800
600
HC
AMS
SMS
D
Density
1.0
0.5
0.0
1e+02
1e+03
1e+04
1e+05
HERV Distance to nearest ZNF gene (bp, log10 scale)
E
chr19

**Supplemental Figure 2. MS-specific HERV expression patterns. (A)** A heatmap comparing the general HERV expression across ERVmancer (right), ERVmap (middle) and TELEscope (left) in long- and short-read data. SMS samples generally show higher HERV expression. **(B)** HERV expression of 11 HERVs found on chr19 with MWU unadjusted p-values significant for greater expression in SMS vs HC and AMS states. **(C)** HERV and clade level expression for 58 HERVs or clades with less than 10 nodes deep from a HERV on chromosome 19, which have an MWU unadjusted significant p-value for greater expression in SMS vs HC and AMS states. **(D)** Density plot showing the log10 distance in base pairs of the HERV to the nearest Zinc Finger (ZNF) gene. **(E)** PhenoGram[4] plot showing the hg19 position on chr19 of the HERVs and clades with significant MWU unadjusted p-values for higher expression in SMS than the other states.

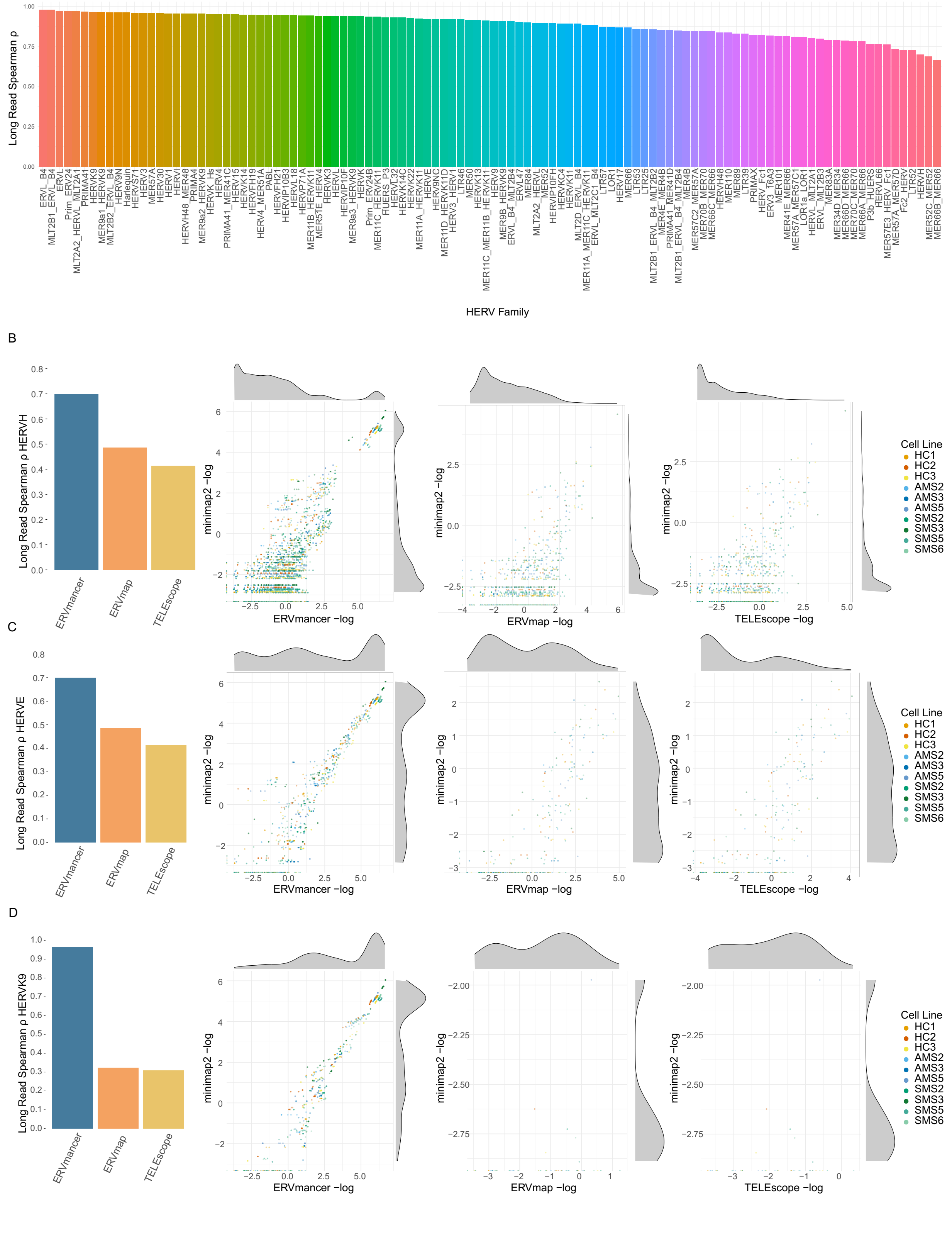


**Supplemental Figure 3. HERV family specific performance evaluation. (A)** ERVmancer correlations with long-read output by HERV family or subtype. **(B)-(D)** Left: Correlation of the short-read quantification method counts with long-read output for the specified HERV family. Right: Scatterplots of the -$\log_2$ normalized counts on the x-axis and the -$\log_2$ normalized long read counts on the y-axis for ERVmancer, ERVmap, and TELEscope. **(B)** HERVH **(C)** HERVE **(D)** HERVK9.

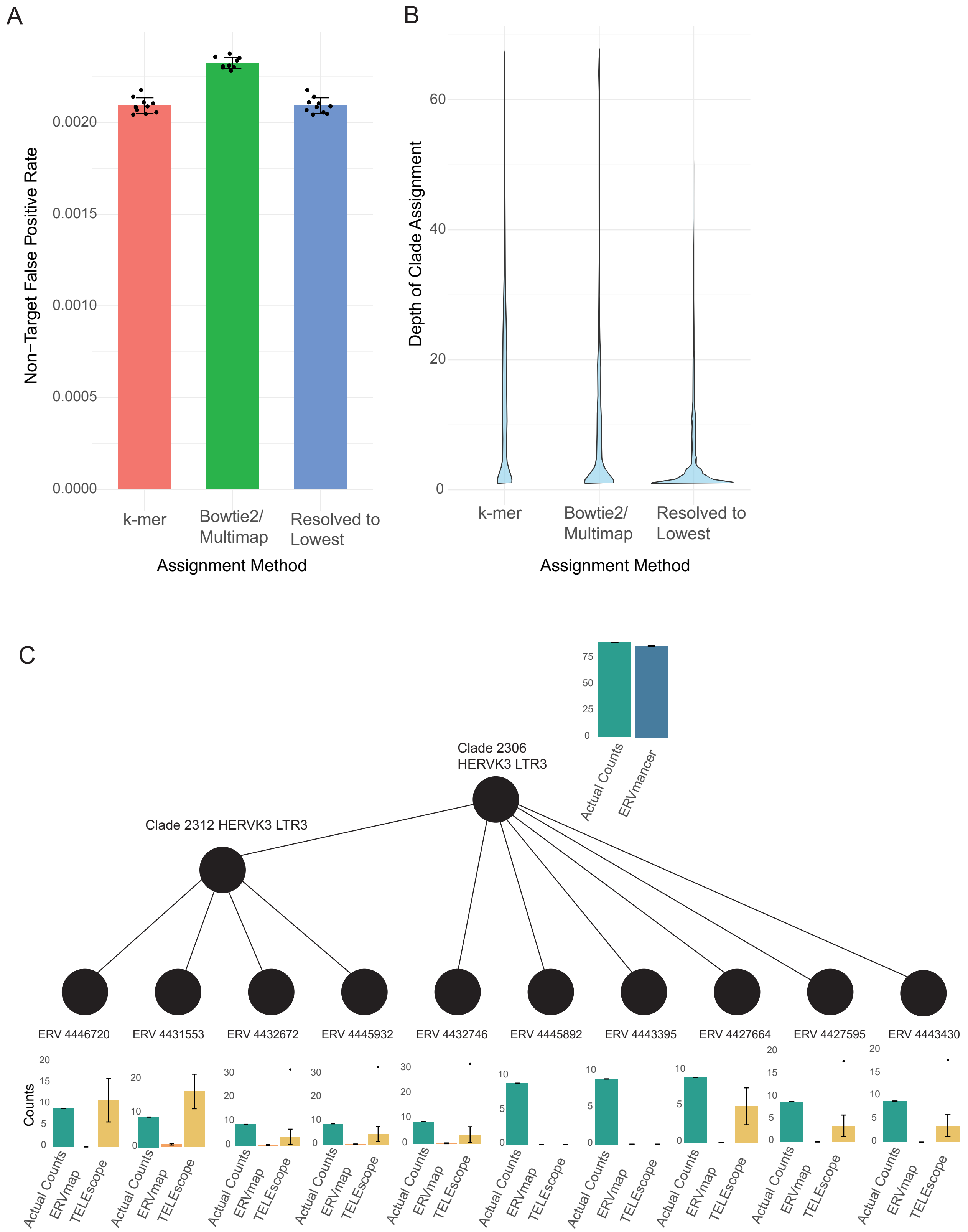


**Supplemental Figure 4. Performance evaluation using simulated sequencing data (A),(B)** Comparison of the k-mer or Bowtie2 multimapping steps of ERVmancer in isolation, versus the final combined approach mapping to the lowest clade across the two methods. **(A)** Non-target false positive rate for the k-mer approach, the Bowtie2 multimapping approach, and the final ERVmancer method. **(B)** The depth of the clade assignment between the k-mer, Bowtie2, and final ERVmancer method. **(C)** Comparison of the different methods in quantifying the HERVs in Clade 2306. ERVmancer assigns reads correctly to Clade 2306, while the other methods either over filter, as in the case of ERVmap, or misassign within the clades, as seen in TELEscope. Some clades are not shown for simplicity.

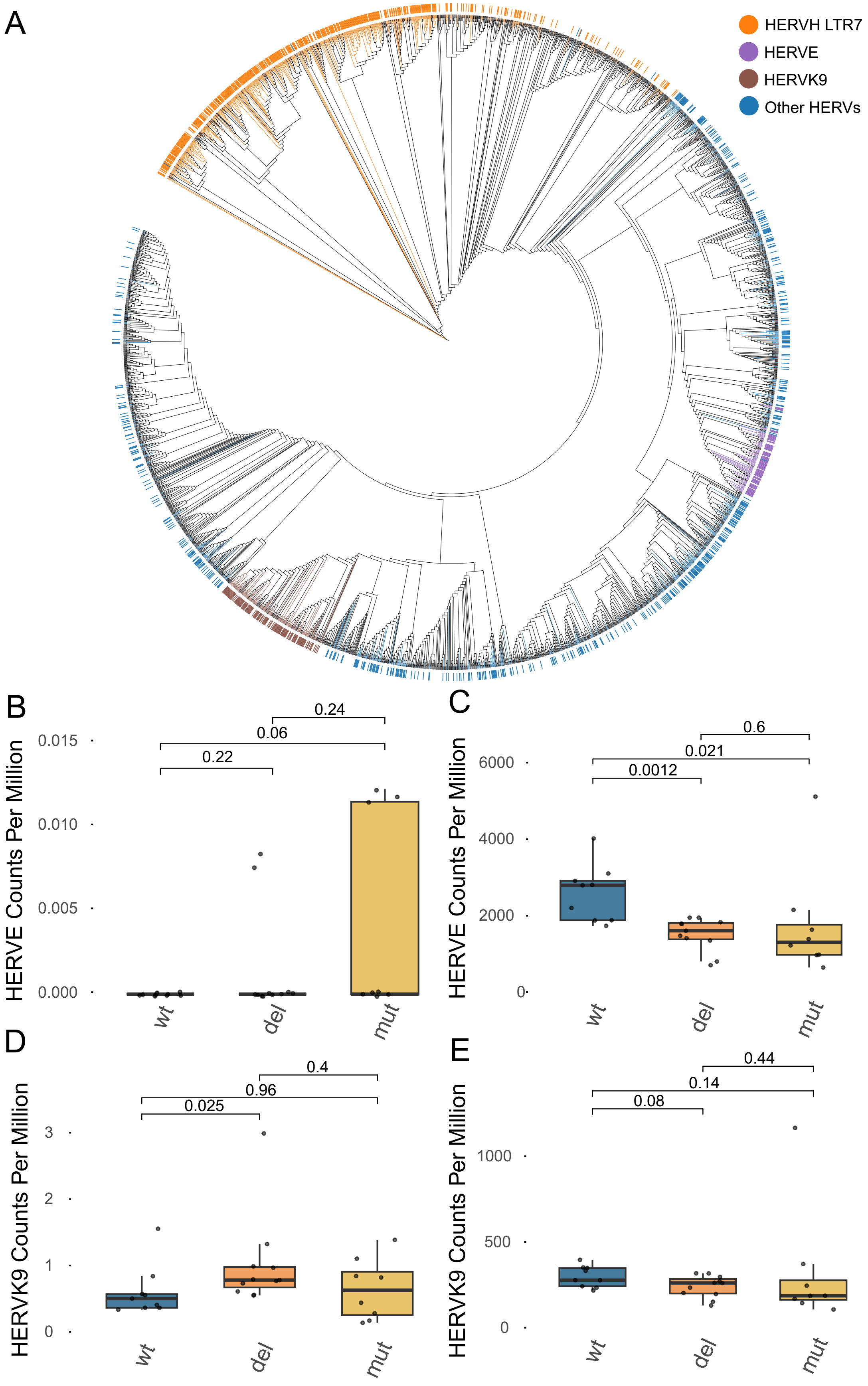
A
HERVH LTR7
HERVE
HERVK9
Other HERVs
B
HERVE Counts Per Million
0.24
0.06
0.22
0.015
0.010
0.005
0.000
wt
del
mut
C
HERVE Counts Per Million
0.6
0.021
0.0012
6000
4000
2000
0
wt
del
mut
D
HERVK9 Counts Per Million
0.4
0.96
0.025
3
2
1
0
wt
del
mut
E
HERVK9 Counts Per Million
0.44
0.14
0.08
1000
500
0
wt
del
mut

**Supplemental Figure 5. p53 binding sites across diverse HERV clades and families. (A)** The phylogenetic tree colored by different HERV clades with p53 binding sites. **(B),(C)** HERVE CPM for HERVE with p53 binding sites which are individually evaluating expression greater (B) or less than (C) the WT samples (MWU unadjusted p-values are reported). **(D),(E)** HERVK9 CPM for HERVK9 with p53 binding sites which are individually evaluated for expression greater (D) or less than (E) the WT samples (MWU unadjusted p-values are reported).
